# A conserved amino acid in the C-terminus of HPV E7 mediates binding to PTPN14 and repression of epithelial differentiation

**DOI:** 10.1101/2020.05.22.111740

**Authors:** Joshua Hatterschide, Alexis C. Brantly, Miranda Grace, Karl Munger, Elizabeth A. White

**Affiliations:** Department of Otorhinolaryngology: Head and Neck Surgery, University of Pennsylvania Perelman School of Medicine, Philadelphia, PA, USA; Department of Developmental, Molecular and Chemical Biology, Tufts University School of Medicine, Boston, MA, USA

**Author notes:** Correspondence: Elizabeth A. White.

## Abstract

The human papillomavirus (HPV) E7 oncoprotein is a primary driver of HPV-mediated carcinogenesis. The E7 proteins from diverse HPV bind to the host cellular non-receptor protein tyrosine phosphatase type 14 (PTPN14) and direct it for degradation through the activity of the E7-associated host E3 ubiquitin ligase UBR4. Herein we show that a highly conserved arginine residue in the C-terminal domain of diverse HPV E7 mediates interaction with PTPN14. We found that disruption of PTPN14 binding through mutation of the C-terminal arginine did not impact the ability of several high-risk HPV E7 proteins to bind and degrade the retinoblastoma tumor suppressor or activate E2F target gene expression. HPVs infect human keratinocytes and we previously reported that both PTPN14 degradation by HPV16 E7 and PTPN14 CRISPR knockout repress keratinocyte differentiation-related genes. Now we have found that blocking PTPN14 binding through mutation of the conserved C-terminal arginine rendered both HPV16 and HPV18 E7 unable to repress differentiation-related gene expression. We then confirmed that the HPV18 E7 variant that could not bind PTPN14 was also impaired in repressing differentiation when expressed from the complete HPV18 genome. Finally, we found that the ability of HPV18 E7 to extend the lifespan of primary human keratinocytes required PTPN14 binding. CRISPR/Cas9 knockout of PTPN14 rescued keratinocyte lifespan extension in the presence of the PTPN14 binding-deficient HPV18 E7 variant. These results support the model that PTPN14 degradation by high-risk HPV E7 leads to repression of differentiation and contributes to its carcinogenic activity.

**IMPORTANCE:** Human papillomavirus (HPV)-positive carcinomas account for nearly 5% of the global human cancer burden. The E7 oncoprotein is a primary driver of HPV-mediated carcinogenesis. HPV E7 binds and degrades the putative tumor suppressor, PTPN14. However, the impact of PTPN14 binding by E7 on cellular processes is not well defined. Here, we show that PTPN14 binding is mediated by a conserved C-terminal arginine residue of HPV E7 *in vivo*. Additionally, we found that PTPN14 binding contributes to the carcinogenic activity of HPV18 E7 (the second most abundant HPV type in cancers). Finally, we determined that PTPN14 binding by HPV16 E7 and HPV18 E7 represses keratinocyte differentiation. HPV-positive cancers are frequently poorly differentiated and the HPV life cycle is dependent upon the keratinocyte differentiation program. The finding that PTPN14 binding by HPV E7 impairs differentiation has significant implications for both HPV-mediated carcinogenesis and the HPV life cycle.

## INTRODUCTION

Infection with oncogenic human papillomaviruses (HPVs) can cause several epithelial carcinomas including essentially all cervical cancers, a subset of oropharyngeal cancers, and several other anogenital cancers. These HPV-positive cancers account for approximately 5% of the human cancer burden (1). HPVs are non-enveloped dsDNA viruses that infect the basal keratinocytes of cutaneous or mucosal stratified squamous epithelia and couple their replication to keratinocyte differentiation (2). Hundreds of genetically diverse HPV genotypes have been identified to date (3). Only 13-15 of the mucosal-tropic HPVs, termed the ‘high-risk’ HPVs, are significantly associated with carcinogenesis. Infection with a high-risk HPV is the first step in the carcinogenic process. Cells persistently infected with high-risk HPV can accumulate secondary hits such as viral genome integration into a host chromosome. This can lead to dysregulated, constitutive expression of the viral E6 and E7 oncoproteins, which are required for keratinocyte immortalization and for the transformed phenotype in cervical carcinoma cell lines (4–8).

HPV E6 and E7 are multifunctional viral proteins that lack enzymatic activity. They can alter host cellular processes through interactions with cellular proteins. One activity of many HPV E7 is to bind and inactivate the retinoblastoma tumor suppressor (RB1) (9, 10). This results in the release of E2F transcription factors and promotes entry into the cell cycle. High-risk HPV E7 additionally promote the proteasomal degradation of RB1, in the case of HPV16 E7 through the recruitment of a Cullin 2-Zer1 E3 ubiquitin ligase complex (11, 12). High-risk HPV E6 direct the tumor suppressor p53 for proteasome-mediated degradation (13, 14), blocking the induction of apoptosis that would arise from E7-mediated stabilization of p53 (15, 16).

Binding to RB1 is necessary but not sufficient to explain the carcinogenic activities of HPV E7 (17–27). We have begun to characterize a second putative tumor suppressor, the non-receptor protein tyrosine phosphatase PTPN14, that is bound and degraded by HPV E7 (11, 28, 29). High-risk HPV E7 direct PTPN14 for proteasome-mediated degradation dependent on the interaction between E7 and the host E3 ubiquitin ligase UBR4 (28, 30). Our early experiments determined that amino terminal sequences in HPV E7 were required for the interaction with UBR4 whereas binding to PTPN14 mapped broadly to the C-terminus of E7. HPV E7 binds to the conserved C-terminal protein tyrosine phosphatase domain of PTPN14. The E7/UBR4/PTPN14 complex is genetically and biochemically separable from the E7/RB1 complex (28, 29).

PTPN14 is proposed to inhibit the HIPPO pathway transcriptional coactivators YAP1 and TAZ. The inhibitory activity of PTPN14 may involve interaction(s) with the HIPPO pathway regulators KIBRA and LATS1 (31–33) and/or direct binding of PTPN14 to YAP1, promoting YAP1 cytoplasmic sequestration (34). PTPN14 phosphatase activity is not required for its inhibition of YAP1. The consequences of PTPN14 degradation in HPV E7-expressing or HPV-positive cells are incompletely understood.

To determine the effect of PTPN14 degradation by HPV E7, we have begun to characterize E7 variants that cannot degrade PTPN14. We recently described a variant of HPV16 E7, HPV16 E7 E10K, that was found among HPV16 sequences from clinical samples (35). HPV16 E7 E10K could not bind to UBR4 and consequently could not direct PTPN14 for degradation (29). Our experiments indicated that PTPN14 degradation by HPV16 E7 impaired the expression of differentiation-related genes in keratinocytes. Our data also supported that PTPN14 degradation could contribute to the carcinogenic activity of high-risk HPV E7. Compared to prototypical HPV16 E7, HPV16 E7 E10K was impaired in its ability to immortalize HFKs when co-expressed with HPV16 E6.

Yun and colleagues recently reported the crystal structure of the phosphatase domain of PTPN14 in complex with the C-terminal domain of HPV18 E7 (36). The two proteins bind directly and with high affinity (*K*_D_=18.2 nM). Several interactions between amino acids on PTPN14 and on HPV18 E7 contribute to the protein-protein interaction, including an electrostatic interaction of HPV18 E7 Arg84 with PTPN14 Glu1095. *In vitro* binding experiments confirmed our previous observation that E7 proteins encoded by diverse HPV bind to PTPN14. Using a structure-guided mutational approach, the authors designed a PTPN14-binding deficient HPV18 E7 and found that it was impaired in promoting keratinocyte growth and migration. The C-terminal arginine that was identified by Yun and colleagues to directly contribute to PTPN14 binding is highly conserved among HPV E7. Eighty-three percent of genus alpha E7 have an arginine at this position and 92% have a positively charged residue.

Here we show that the conserved C-terminal arginine is required for diverse HPV E7 proteins to bind and degrade PTPN14 in cells. Consistent with our previous results, we find that PTPN14 binding contributes to the ability of high-risk HPV E7 to repress keratinocyte differentiation. We extend this result to include cells that contain the complete HPV genome. Finally, we find that PTPN14 binding by HPV18 E7 is required for the keratinocyte lifespan extension activity of HPV18 E7.

## RESULTS

### Mutation of a conserved C-terminal arginine residue in diverse HPV E7 restores PTPN14 protein levels

HPV E7 proteins form a complex with PTPN14 and the host E3 ubiquitin ligase UBR4 to direct PTPN14 for proteasome mediated degradation (28). We have shown that mutations in HPV16 E7 that abrogate UBR4 binding also impair degradation of PTPN14 and prevent HPV16 E7 from repressing differentiation gene expression (29). However, we had not characterized any variant of E7 that did not bind to PTPN14 nor had we extended our analysis to include genotypes other than HPV16.

HPV E7 proteins from diverse genotypes have some conserved and some divergent amino acid sequences (Figure 1A). Diverse HPV E7 proteins bind to PTPN14 (11, 28). Given the connection we recently established between PTPN14 and differentiation (29), we re-examined the relationship between diverse genus alpha HPV E7 and PTPN14 protein levels. We transduced hTert-immortalized human foreskin keratinocyte (N/Tert-1) cells with retroviral vectors encoding HPV E7 from several genus alpha HPV types. Using newly derived, passage-matched keratinocyte pools stably expressing HPV E7, we found that E7 proteins from high-risk, low-risk, and non-cancerous genus alpha HPV reduced steady-state PTPN14 protein levels (Figure 1B). Consistent with our previous results, high-risk HPV18 E7 expression resulted in the most substantial reduction in PTPN14 protein levels.

**Figure 1.**
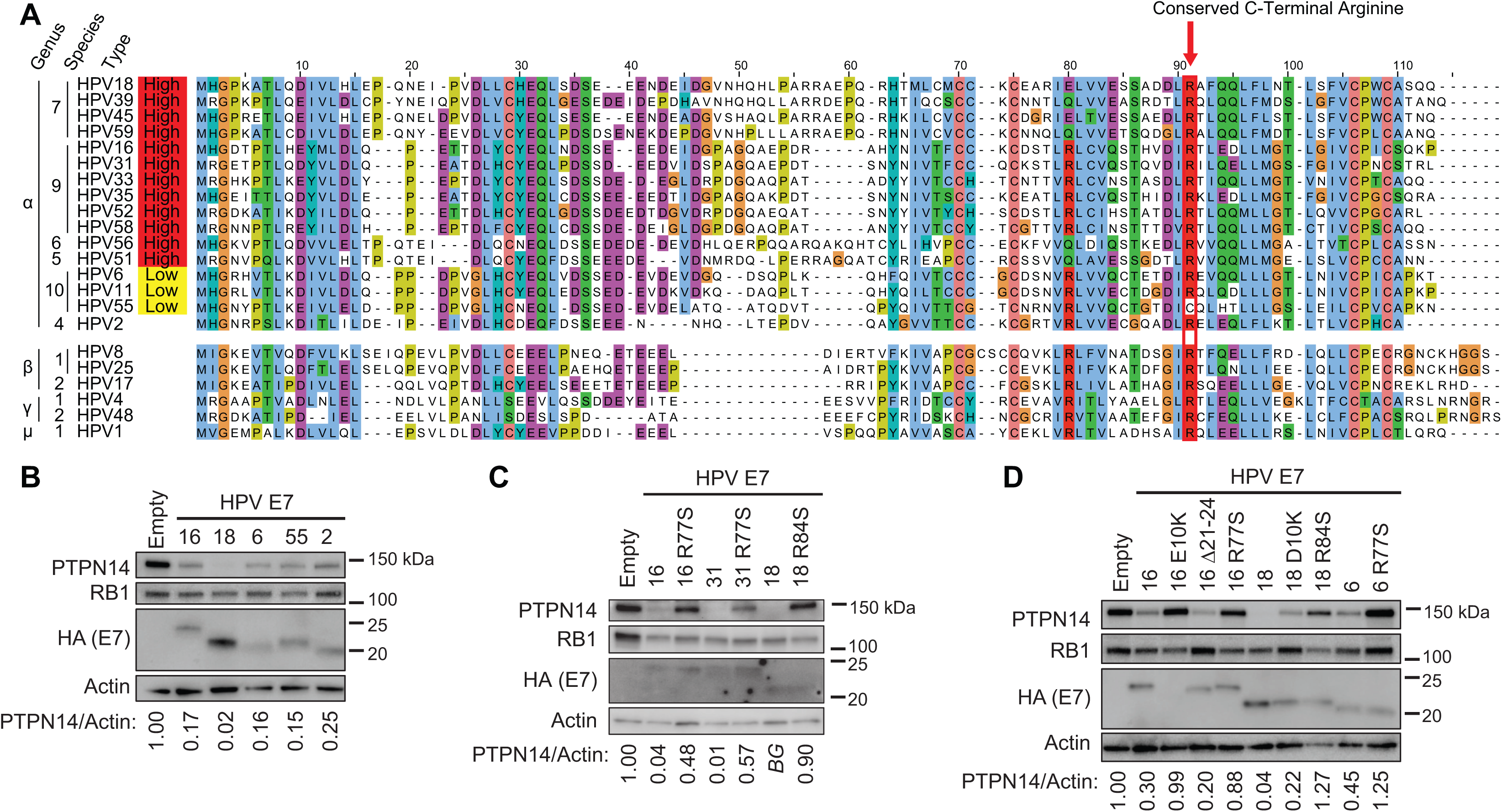
Conserved PTPN14 binding and degradation by HPV E7. (A) Protein sequence alignment of 22 HPV E7 ORFs from diverse HPV genotypes. Alignments were generated using the Clustal Omega algorithm and colored using the ClustalX color scheme. (B) Cell lysates from N/Tert-1 cells expressing E7 proteins from various HPV genotypes were subjected to SDS/PAGE/Western analysis and probed with antibodies to PTPN14, RB1, HA (E7) and Actin. (C and D) Cell lysates from N/Tert-1 cells expressing WT or variant HPV16, HPV31, HPV18, and HPV6 E7 were subjected to SDS/PAGE/Western analysis and probed with antibodies to PTPN14, RB1, HA (E7) and Actin. Band intensity was quantified using ImageJ. Values represent PTPN14 band intensity relative to Actin band intensity. *BG* indicates that band intensity was ≤ background intensity.

We and others previously determined that PTPN14 binding maps broadly to the carboxyl terminus of HPV E7 (28, 30). A recent crystal structure of the C-terminus of HPV18 E7 in complex with the PTP domain of PTPN14 determined that the two proteins bind directly and elucidated the PTPN14-HPV18 E7 binding interface (36). Several residues of HPV18 E7 were found to be critical for this interaction, including Arg84 (R84; R77 in HPV16 E7) which makes an electrostatic interaction with Glu1095 of PTPN14. The corresponding arginine residue is highly conserved among HPV E7 proteins. It is completely conserved among the prototypical E7 open reading frames (ORFs) from high-risk HPV (Figure 1A), 83% conserved among E7 ORFs from genus alpha, and 80% conserved among E7 ORFs from 193 diverse HPV genotypes. The high degree of conservation of the C-terminal arginine may indicate strong selective pressure for HPV E7 to maintain the ability to bind PTPN14.

The requirement for the C-terminus of E7 to bind PTPN14 led us to characterize a naturally occurring variant of HPV16 E7 identified by Mirabello and colleagues (35). The variant, HPV16 E7 R77S, has a mutation in the conserved arginine residue (Arg77 to Ser; R77S). We hypothesized that this HPV16 E7 might be unable to bind and degrade PTPN14. To test if mutation of the C-terminal arginine impairs PTPN14 degradation by HPV E7 proteins, we engineered the arginine to serine mutation into the E7 ORFs from HPV16, HPV31, HPV18, and HPV6. We then transduced N/Tert-1 cells with retroviruses encoding the E7 variants, the corresponding wild-type/prototypical (WT) E7s, and several additional mutants as controls. The control variants are: HPV16 E7 E10K (does not bind UBR4) (29), HPV16 E7 Δ21-24 (does not bind RB1) (10), and HPV18 E7 D10K (contains the mutation analogous to HPV16 E7 E10K). We found that cells expressing HPV16 E7 R77S, HPV31 E7 R77S, HPV18 E7 R84S, and HPV6 E7 R77S exhibited higher PTPN14 protein levels than the corresponding WT E7-expressing cell lines (Figure 1C and 1D). This is consistent with the idea that the C-terminal arginine is required for diverse genus alpha HPV E7 proteins to interact with PTPN14. In agreement with our previous results, PTPN14 expression was restored in cells expressing the HPV16 E7 E10K variant (29). HPV18 E7 D10K only partially restored PTPN14 expression (Figure 1D). These results support the notion that the C-terminal HPV E7 binding interface may be highly conserved to mediate PTPN14 binding.

### Mutation of the C-terminal arginine in high-risk HPV E7 does not affect RB1 degradation or E2F target gene activation

In addition to binding PTPN14, HPV E7 proteins bind and inactivate RB1 (9, 10). High-risk HPV E7 proteins additionally direct RB1 for proteasomal degradation (11, 12). We found that HPV16 E7 R77S, HPV31 E7 R77S, and HPV18 E7 R84S were comparable to WT E7 proteins with respect to their capacity to reduce RB1 steady-state protein levels (Figure 1C). To test whether HPV16 E7 R77S and HPV18 E7 R84S could induce the expression of E2F target genes downstream of RB1 inactivation, we transduced primary human foreskin keratinocytes (HFK) with retroviral vectors encoding a panel of HPV16 and HPV18 E7 variants. Total RNA was collected from these cell lines and the expression of the E2F target genes *CCNE2* and *MCM2* was assessed by qRT-PCR. With the exception of HPV16 E7 Δ21-24, which cannot bind RB1, all HPV16 and HPV18 E7 variants including HPV16 E7 R77S and HPV18 E7 R84S were able to induce E2F target gene expression (Figure 2A and B). We conclude that mutation of the C-terminal arginine does not affect RB1 inactivation by high-risk HPV E7 proteins.

**Figure 2.**
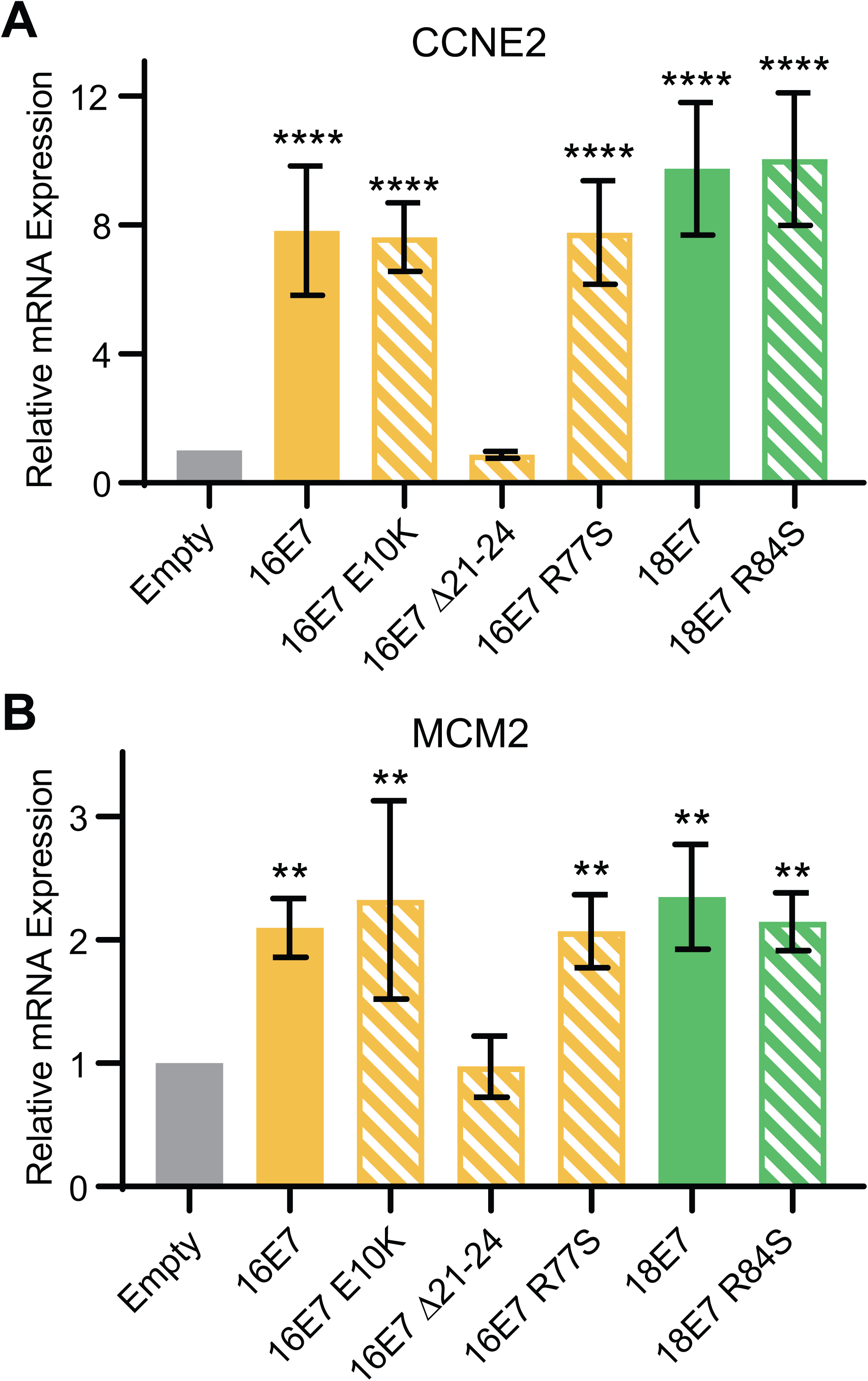
PTPN14 binding and degradation by HPV E7 are not required to promote the expression of E2F-regulated genes. Primary HFK were transduced with retroviral vectors encoding various HPV16 and HPV18 E7 proteins. qRT-PCR was used to measure the expression of the E2F-regulated genes (A) *CCNE2* and (B) *MCM2* relative to *GAPDH*. Graphs display the mean ± SD of three independent replicates. Statistical significance was determined by ANOVA (\*\**p* < 0.01, \*\*\*\**p* < 0.0001).

### Mutation of the conserved C-terminal arginine disrupts PTPN14 binding by high-risk HPV E7

To confirm that mutation of the conserved C-terminal arginine disrupted PTPN14 binding, we immunoprecipitated E7 from the N/Tert cell lines that stably express Flag and HA-tagged HPV16 and HPV18 E7 and variant E7. We found that PTPN14 did not coimmunoprecipitate with HPV16 E7 R77S or HPV18 E7 R84S (Figures 3A and 3B). In contrast PTPN14 coimmunoprecipitated with HPV16 E7 WT, HPV18 E7 WT, and HPV16 E7 Δ21-24, which cannot bind RB1. In agreement with the analysis of total cell protein and RNA (Figures 1 and 2), HPV16 E7 R77S and HPV18 E7 R84S input cell lysates contained more PTPN14 protein than the respective WT E7 cell lysates. HPV16 E7 R77S and HPV18 E7 R84S retained the capacity to coimmunoprecipitate with RB1 and reduce steady state RB1 levels. These data support the model that diverse HPV E7 require the conserved C-terminal arginine to bind to PTPN14 in keratinocytes.

**Figure 3.**
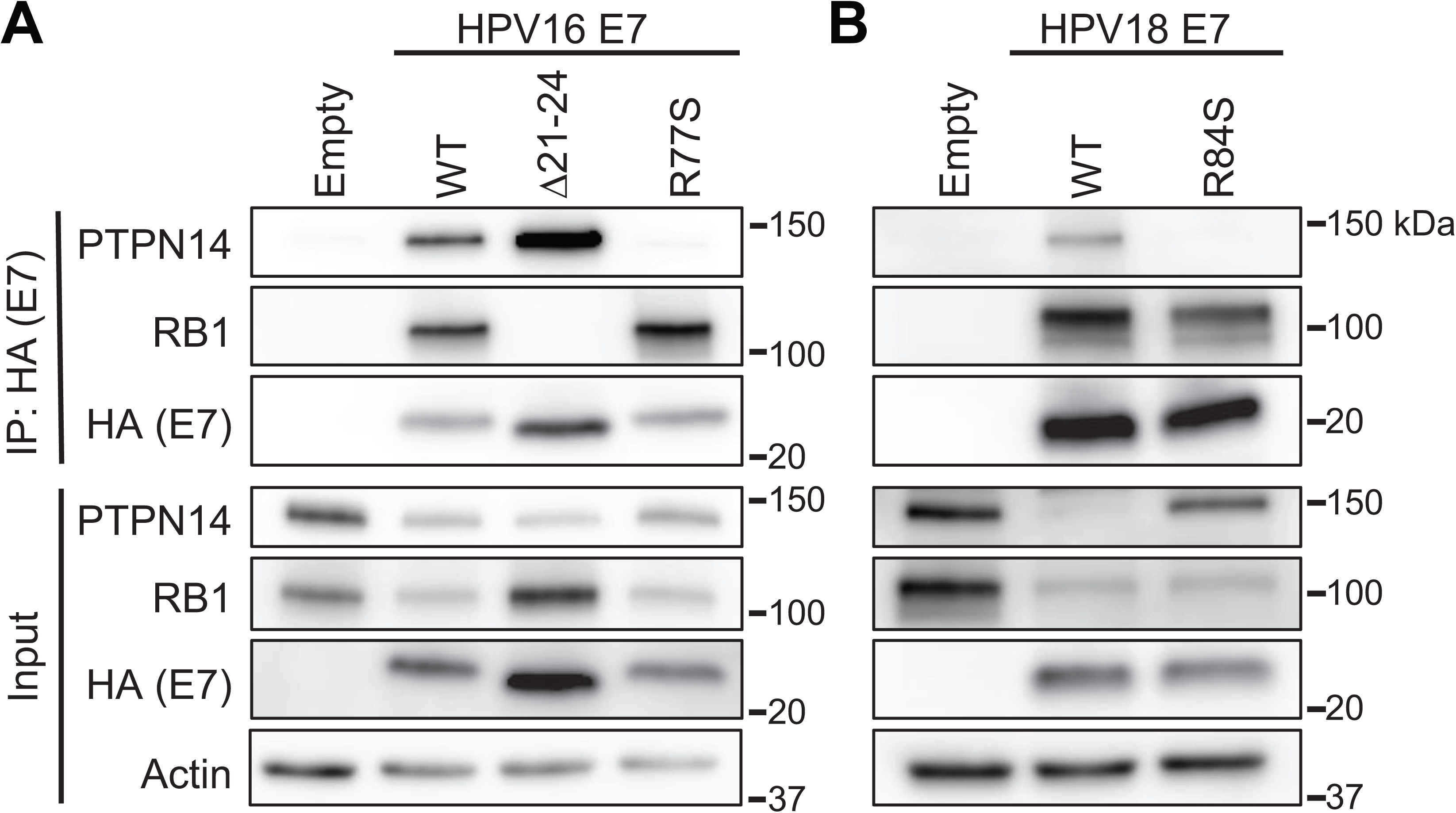
HPV16 E7 R77S and HPV18 E7 R84S cannot bind to PTPN14. N/Tert-1 keratinocytes were transduced with retroviruses encoding various HPV16 E7 (A), HPV18 E7 (B), or the empty vector control. HPV E7-FlagHA was immunoprecipitated with anti-HA beads from total cell lysates and coimmunoprecipitation of PTPN14 and RB1 was assessed by SDS/PAGE/Western analysis (upper). Total cell lysates were subjected to SDS/PAGE/Western analysis and probed with antibodies to PTPN14, RB1, HA (E7) and actin (lower).

### PTPN14 binding is required for high-risk HPV E7 to repress differentiation

We previously determined that PTPN14 degradation by HPV16 E7 is required for an E7-mediated repression of keratinocyte differentiation (29). Therefore, we hypothesized that impairing PTPN14 binding by mutating the C-terminal arginine residue would likewise render high-risk HPV E7 proteins unable to repress differentiation. We used a panel of N/Tert cell lines expressing WT and arginine to serine mutated HPV E7 proteins from the high-risk HPV16 and HPV18. Cells were induced to differentiate by culture in growth factor-free media supplemented with 1.5 mM CaCl_2_. We collected whole cell lysates at 0 and 72 h post-differentiation and analyzed cytokeratin 1 (KRT1), cytokeratin 10 (KRT10), and involucrin (IVL) by Western blot. KRT1 and KRT10 are early differentiation markers and IVL is a component of the cornified envelope. We found that HPV16 E7 WT and HPV18 E7 WT were able to repress the expression of all three differentiation markers, whereas HPV16 E7 R77S and HPV18 E7 R84S were not (Figure 4).

**Figure 4.**
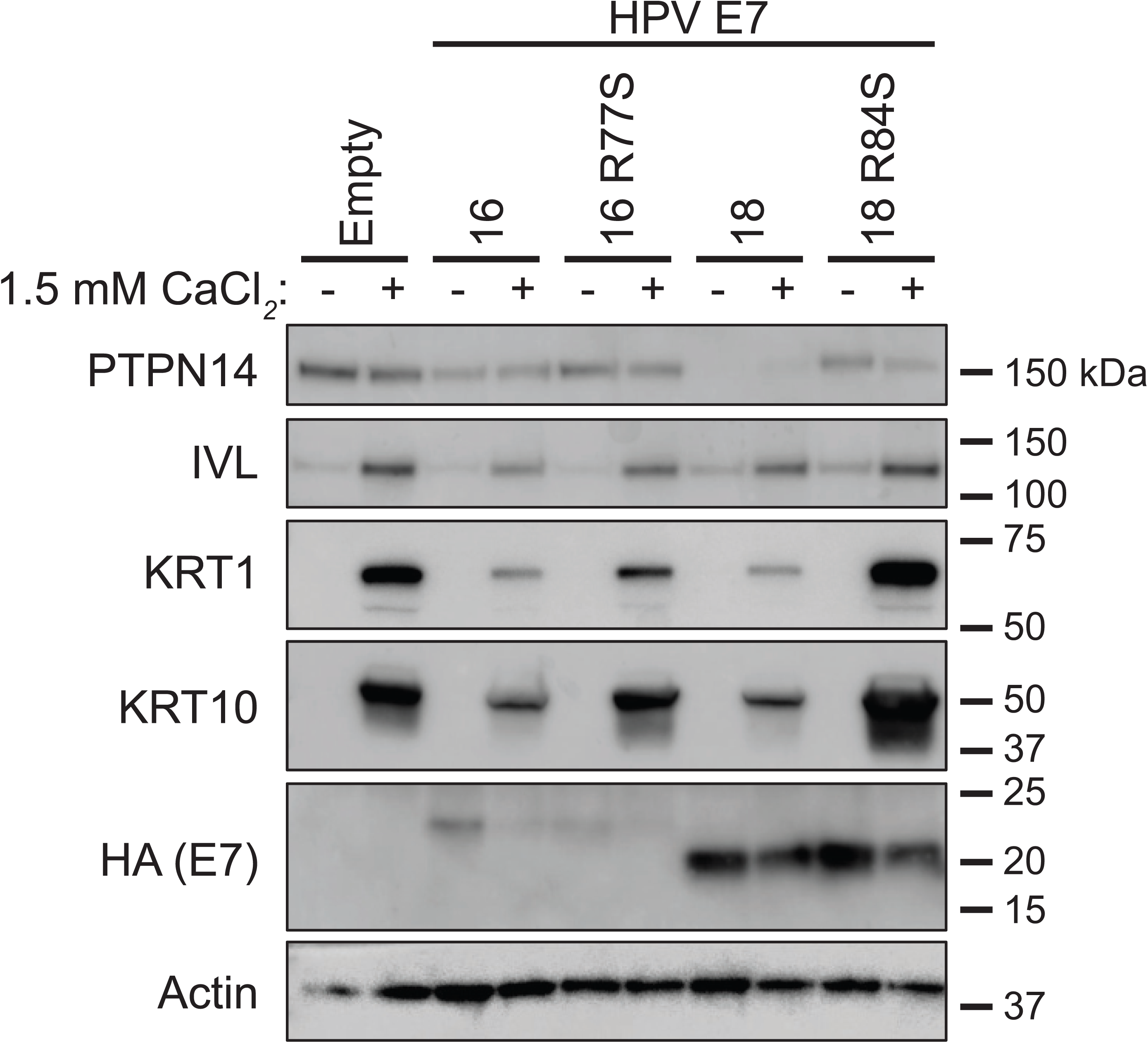
HPV16 E7 R77S and HPV18 E7 R84S are impaired in their ability to repress differentiation gene expression induced by calcium treatment. N/Tert-1 keratinocytes expressing HPV16 E7 WT and R77S, HPV18 E7 WT and R84S or the empty vector control were maintained in growth medium (-) or were induced to differentiate with 1.5 mM CaCl_2_ for 72h (+). Cell lysates were subjected to SDS/PAGE/Western analysis and probed with antibodies to PTPN14, IVL, KRT1, KRT10, HA (E7) and actin.

Because HPV18 E7 leads to the greatest depletion in PTPN14 protein levels, we chose to focus on HPV18 E7 for the remainder of this study. Loss of contact with the basement membrane induces keratinocyte differentiation *in vivo*. This can be simulated in cultured cells by growing keratinocytes in suspension (37–39), which tends to promote high expression of markers of late differentiation. To test whether PTPN14 binding by HPV E7 also impairs differentiation in response to detachment, we cultured our N/Tert cell lines expressing HPV18 E7 WT, HPV18 E7 R84S, and the empty vector control in ultra-low adherence plates for 12 h. We collected RNA from adherent cells or cells in suspension and measured the expression of the late differentiation marker *IVL* by qRT-PCR. *IVL* mRNA expression was induced in all cell lines after suspension. However, HPV18 E7 R84S displayed increased *IVL* expression post-differentiation compared to HPV18 E7 WT (Figure 5A), mirroring our results obtained from CaCl_2_ differentiation. These data indicate that PTPN14 binding by HPV18 E7 is necessary for the repression of differentiation gene expression following differentiation stimuli.

**Figure 5.**
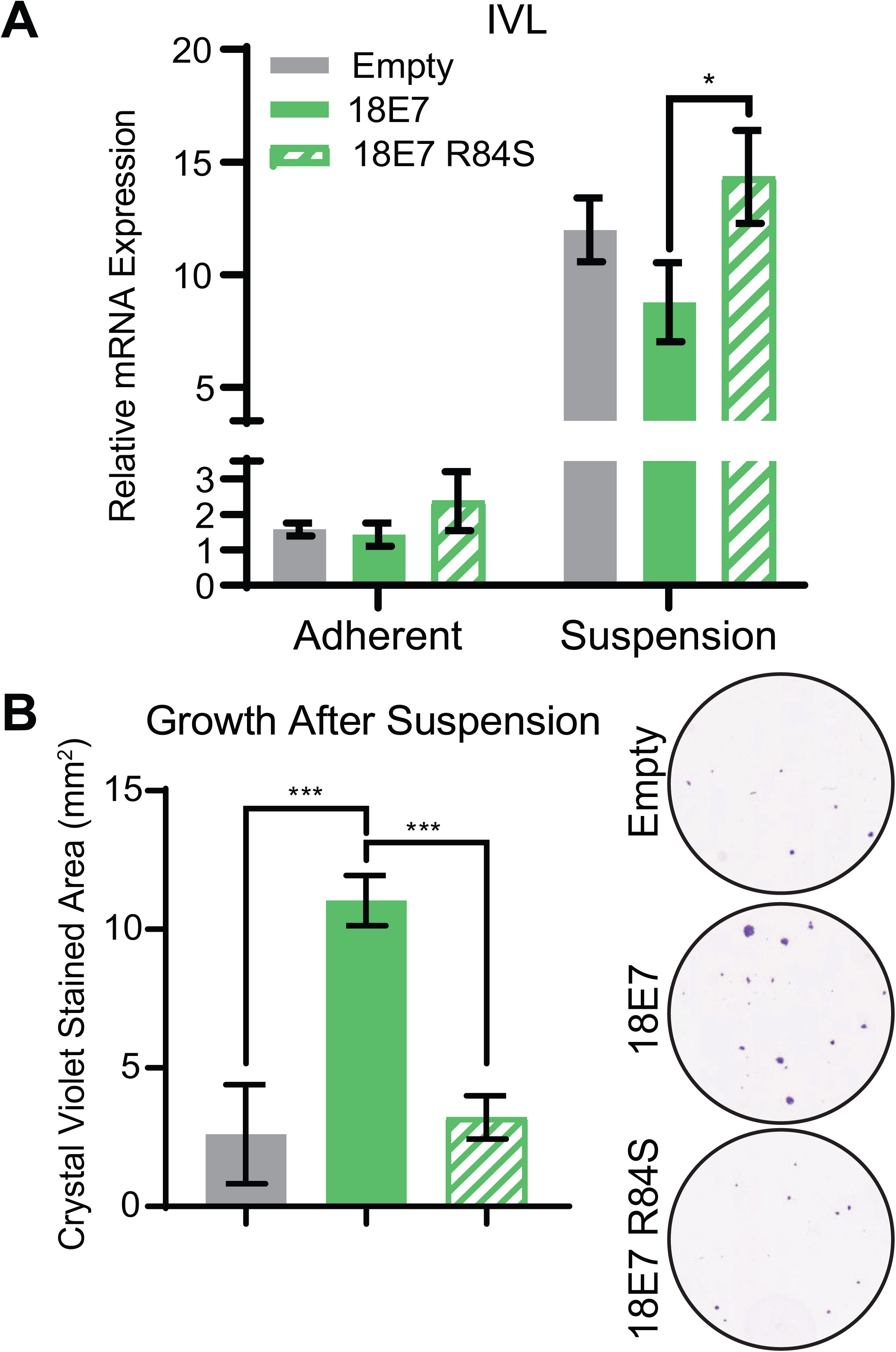
HPV18 E7 binding to PTPN14 contributes to the repression of differentiation induced by suspension and improves survival of cells grown in suspension. N/Tert-1 keratinocytes expressing HPV18 E7 WT, HPV18 E7 R84S, or the empty vector control were induced to differentiate by culturing in ultra-low adherence plates for 12h. (A) qRT-PCR was used to measure the expression of the differentiation marker IVL relative to GAPDH in adherent or suspended cells. Graph displays the mean ± SD of three independent replicates. (B) Survival after suspension was assessed by replating 1,000 cells from suspension, allowing cells to grow for 5 days, and measuring total viable cell area by crystal violet staining. Graph displays the mean ± SD of three independent replicates. Representative images of crystal violet staining are displayed. Statistical significance was determined by ANOVA (\**p* < 0.05, \*\*\**p* < 0.001).

Keratinocytes exit the cell cycle after differentiating, which leads to decreased growth capacity. To test if PTPN14 binding by HPV18 E7 affects cellular viability after suspension, we re-plated 1,000 cells that were grown in ultralow adherence plates for 12 h onto standard tissue culture plates and cultured them for an additional 5 days. We then fixed and stained them with crystal violet and measured the area covered by viable cells. We found that HPV18 E7 WT increased viability after stimulus to differentiate in suspension, while the growth of HPV18 E7 R84S expressing cells resembled the empty vector control (Figure 5B). These data support that PTPN14 binding by high-risk HPV E7 is required for the impairment of keratinocyte differentiation, promoting cellular viability following a differentiation stimulus.

### Repression of differentiation is the predominant effect of PTPN14 binding by HPV18 E7 on the transcriptome of primary keratinocytes

We hypothesized that comparing HPV18 E7 and HPV18 E7 R84S in an unbiased analysis of gene expression might reveal additional cellular processes that are up- or down-regulated by E7-mediated PTPN14 binding and degradation. To test this, we transduced primary HFKs with retroviral vectors encoding HPV18 E7 WT, HPV18 E7 R84S, or the empty vector control. Triplicate primary cell populations were established and selected with puromycin. RNA isolated from these cell populations was polyA-selected and analyzed by RNA-sequencing (RNA-seq). We found that 271 genes were differentially regulated between HFK expressing HPV18 E7 WT and HPV18 E7 R84S with cut offs of >1.5-fold change and false discovery rate (FDR) adjusted *p*-value of <0.05. The majority of these genes (247) were found to be more repressed in cells expressing HPV18 E7 WT than those expressing the HPV18 E7 R84S mutant (Figure 6A). We performed a gene ontology (GO) enrichment analysis on the list of genes with lower expression in wild type HPV18 E7 expressing cells and found that this list was highly enriched for GO terms related to keratinocyte differentiation (Figure 6B). Further analysis revealed that a high percentage of these 247 genes were related to the following GO terms: Cornification, 15%; Keratinocyte Differentiation, 17%; Skin Development, 19%; and Epidermis Development, 20%. Among the 24 genes that were upregulated in cells expressing HPV18 E7 WT versus cells expressing HPV18 E7 R84S, there were no statistically significantly enriched GO terms.

**Figure 6.**
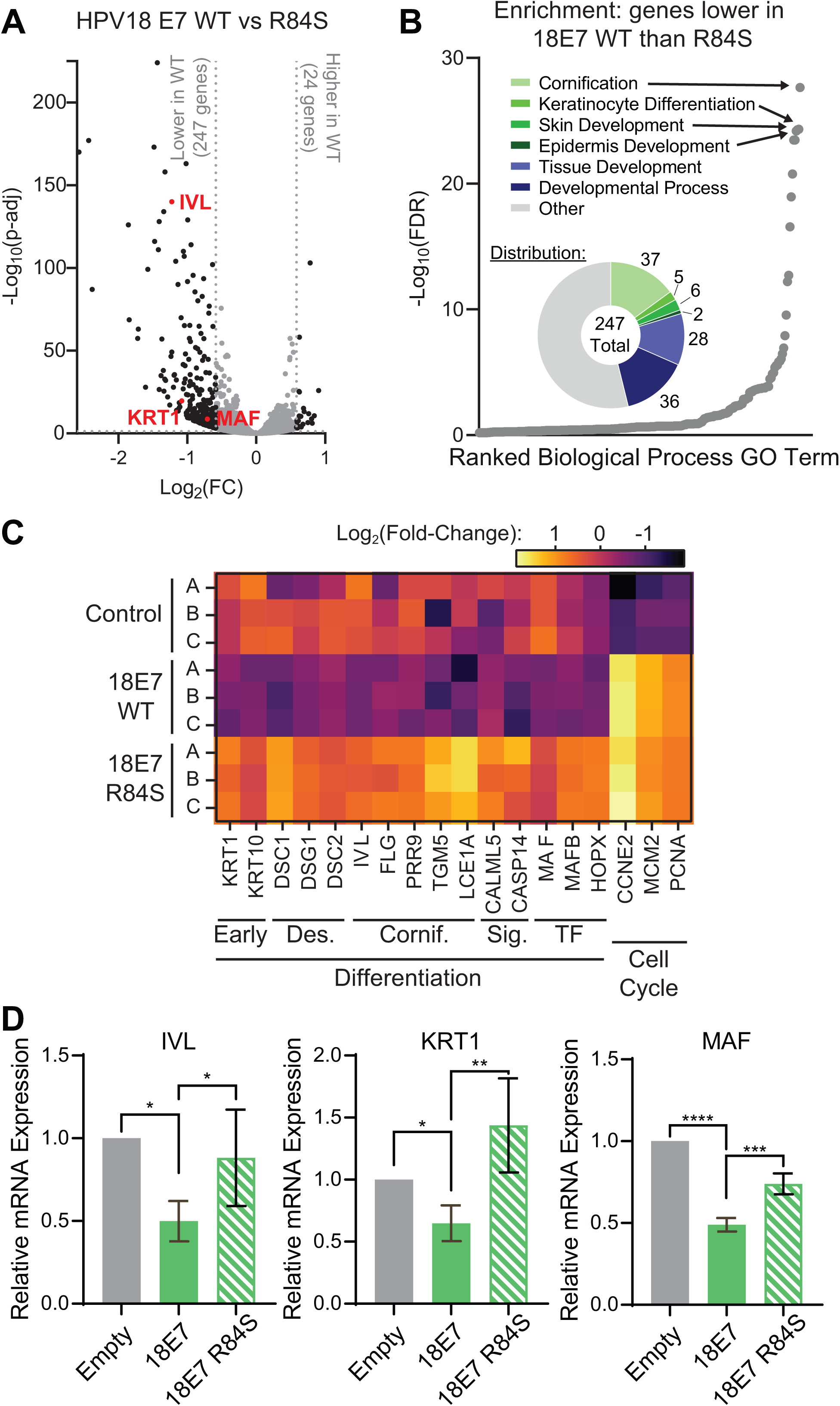
Repression of differentiation is the predominant effect of PTPN14 binding by HPV18 E7 on the transcriptome of primary keratinocytes. Primary HFK were transduced with retroviruses encoding HPV18 E7 WT, HPV18 E7 R84S, or the empty vector control. PolyA-selected RNA collected from early passages was analyzed by RNA-seq. (A) Volcano plot of gene expression changes in HFKs expressing HPV18 WT and HPV18 R84S. Vertical and horizontal dotted lines represent 1.5-fold change and *p*-adj = 0.05, respectively. Representative genes are highlighted in red. (B) Gene Ontology (GO) enrichment analysis of genes down-regulated in HFKs expressing HPV18 WT compared to HPV18 R84S. GO terms are displayed in rank order of their FDR-adjusted *p-*value. Circle chart displays the fraction of down-regulated genes that fall into enriched GO Terms. (C) Heat map displaying the expression patterns of selected genes related to differentiation and the cell cycle. *Early*, early differentiation markers. *Des.*, genes encoding components of the differentiation-associated desmosome complexes. *Cornif.*, genes involved in cornified envelope formation. *Sig.*, genes involved in differentiation-promoting signals. *TF.*, genes encoding differentiation-promoting transcription factors. (D) qRT-PCR was used to measure the expression of the differentiation-related genes *IVL*, *KRT1*, and *MAF* relative to *GAPDH*. Graphs display the mean ± SD of three independent replicates. Statistical significance was determined by ANOVA (\**p* < 0.05, \*\**p* < 0.01, \*\*\**p* < 0.001, \*\*\*\**p* < 0.0001).

Based on the strong differentiation signature we specifically focused on a broad array of differentiation-related genes involved in early differentiation, the differentiation-associated desmosome complexes, cornified envelope formation, differentiation-promoting signaling, and differentiation-promoting transcription factors. Analysis of these selected genes revealed that HPV18 E7 WT indeed broadly represses differentiation-related genes in comparison to the empty vector control and that the expression of these genes is restored in cells that express HPV18 E7 R84S (Figure 6C). Furthermore, this RNA-seq analysis confirmed that both HPV18 E7 WT and HPV18 E7 R84S promote the expression of cell cycle-related genes such as *CCNE2, MCM2*, and *PCNA* that are activated downstream of RB1 degradation. To validate our RNA-seq results, we measured the mRNA expression of several key genes using qRT-PCR. These included *IVL*, *KRT1*, and *MAF*, which is a transcription factor that promotes early differentiation (40). The expression of these three differentiation-related genes was significantly lower in HFK expressing HPV18 E7 WT than those expressing HPV18 E7 R84S or the empty vector control (Figure 6D). In summary, we found that repressing genes related to differentiation was the predominant effect of PTPN14 binding by HPV18 E7 on keratinocyte gene expression in unstimulated keratinocytes.

### The HPV18 E7 C-terminal arginine contributes to differentiation repression in keratinocytes harboring the complete HPV genome

Our data support that PTPN14 binding is required for HPV E7 to repress differentiation when HPV E7 is expressed in the absence of other viral proteins. However, other HPV proteins, particularly E6, have been shown to repress differentiation (41–43). To test whether PTPN14 binding by HPV E7 contributes to repression of differentiation when E7 is expressed from its native context in the complete viral genome, we introduced the HPV18 E7 R84S mutation into pNeo-loxP-HPV18 (44). We transfected the WT and mutant pNeo-loxP-HPV18 vectors along with NLS-Cre into primary HFK and selected stable populations with G418 (Figure 7A). The resulting cell pools are HFK-HPV18^E7 WT^ and HFK-HPV18^E7 R84S^. Uptake and excision of the HPV18 genome were confirmed by PCR from DNA collected at an early passage post-transfection (Figure 7B). To characterize the effect of HPV18 E7 WT and R84S on PTPN14 in the presence of the complete HPV genome, protein lysates were collected from each cell line and compared to lysates from the untransfected parental HFKs. We found that PTPN14 expression was strongly diminished in HFK-HPV18^E7 WT^ but not in HFK-HPV18^E7 R84S^ (Figure 7C). Reduction in RB1 protein levels can serve as a surrogate for high-risk HPV E7 expression. As expected, HFKs harboring either genome displayed reduced RB1 protein levels, suggesting that E7 is expressed from both pNeo-loxP-HPV18 vectors. To test how PTPN14 binding by HPV18 E7 impacts differentiation in the context of the complete genome, these cell populations were stimulated to differentiate with media containing 1.5 mM CaCl_2_. Total RNA was collected 0, 2, and 6 days post differentiation and expression of the early differentiation markers *KRT1* and *KRT10* was measured by qRT-PCR. We found that throughout the time course, HFK-HPV18^E7^ WT exhibited a 2.3- to 5.9-fold reduction in *KRT1* and *KRT10* expression in comparison to HFK-HPV18^E7 R84S^ (Figure 7D). These data indicate that PTPN14 binding by HPV18 E7 contributes to the ability of HPV E7 to repress differentiation gene expression even in the context of the complete HPV genome.

**Figure 7.**
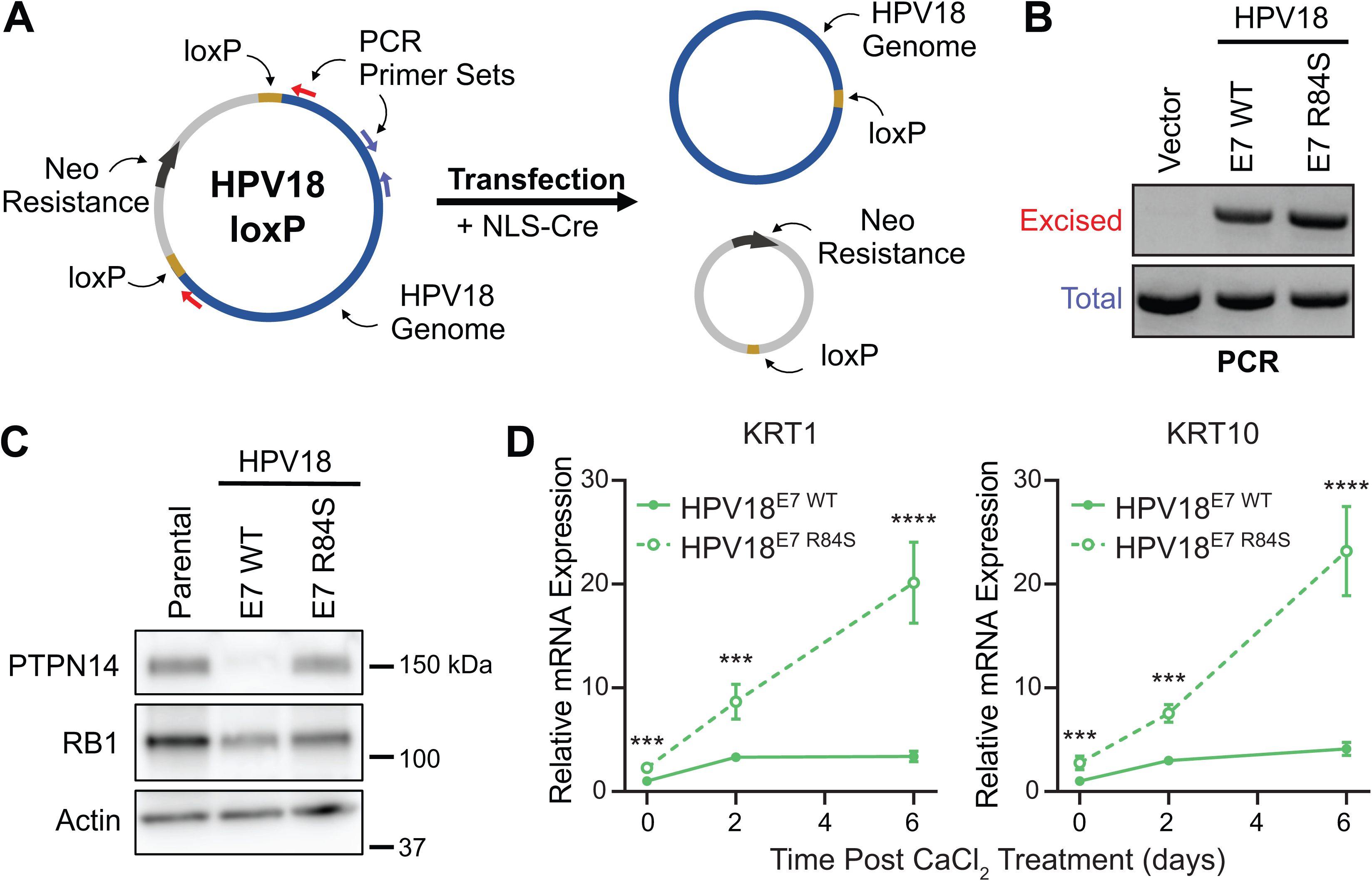
PTPN14 degradation is required for repression of differentiation in primary keratinocytes harboring the HPV18 genome. Primary HFK were transfected with pNeo-loxP-HPV18 vectors and NLS-Cre. (A) Schematic of the pNeo-loxP-HPV18 transfection experimental strategy. (B) PCR/agarose gel analysis of DNA isolated from HFK was used to confirm the transfection and excision of the HPV18 genome following selection. (C) Cell lysates were subjected to SDS/PAGE/Western analysis and probed with antibodies to PTPN14, RB1 and actin. (D) Cells were treated with 1.5 mM CaCl_2_ for 0, 2, and 6 days. qRT-PCR was used to measure the expression of the differentiation markers *KRT1* and *KRT10* relative to *GAPDH*. Graphs display the mean ± SD of two or three independent replicates. Statistical significance was determined by repeated measures two-way ANOVA (\*\*\**p* < 0.001, \*\*\*\**p* < 0.0001).

### PTPN14 binding is required for HPV18 E7 to extend the lifespan of primary human keratinocytes

High-risk HPV E7 extends the lifespan of primary keratinocytes *in vitro* (8, 45). We have previously reported that the E10K mutation in HPV16 E7, which blocks PTPN14 degradation, also impairs its ability to extend the lifespan of primary keratinocytes (29). To assess whether PTPN14 binding by HPV18 E7 is also necessary for lifespan extension by E7, we transduced primary HFK with retroviral vectors encoding HPV18 E7 WT, HPV18 E7 R84S, or the empty vector control and selected for stable populations. By counting cells at each passage, we tracked the number of times each cell population doubled over time. We found that HPV18 E7 R84S was unable to extend the lifespan of primary HFK and that these cells behaved comparably to the HFK-empty vector control (Figure 8A). In contrast, HPV18 E7 WT extended the lifespan of primary HFK until the experiment was terminated 49 days post-transduction. This result supports that PTPN14 binding by HPV18 E7 is essential for its ability to extend the lifespan of primary cells. To establish that lifespan extension by HPV18 E7 is dependent on PTPN14 degradation and not on another activity of HPV18 E7 related to the C-terminal arginine residue, we tested whether deletion of PTPN14 could rescue the defect of the HPV18 E7 R84S mutant and extend keratinocyte life span. These experiments were conducted by co-transducing HFKs from two different keratinocyte donors with the HPV18 E7 retroviral vectors and LentiCRISPR v2 vectors encoding control or PTPN14 targeting guide RNAs (two of each). We found that PTPN14 deletion rescued the defect in the lifespan extension capability of HPV18 E7 R84S to a level that resembled that of HPV18 E7 WT (Figure 8B). PTPN14 KO did not extend the lifespan of cells transduced with the empty vector control and did not further extend the lifespan of cells expressing HPV18 E7 WT. These results further support the model that PTPN14 binding and degradation is required for the carcinogenic activity of high-risk HPV E7 proteins.

**Figure 8.**
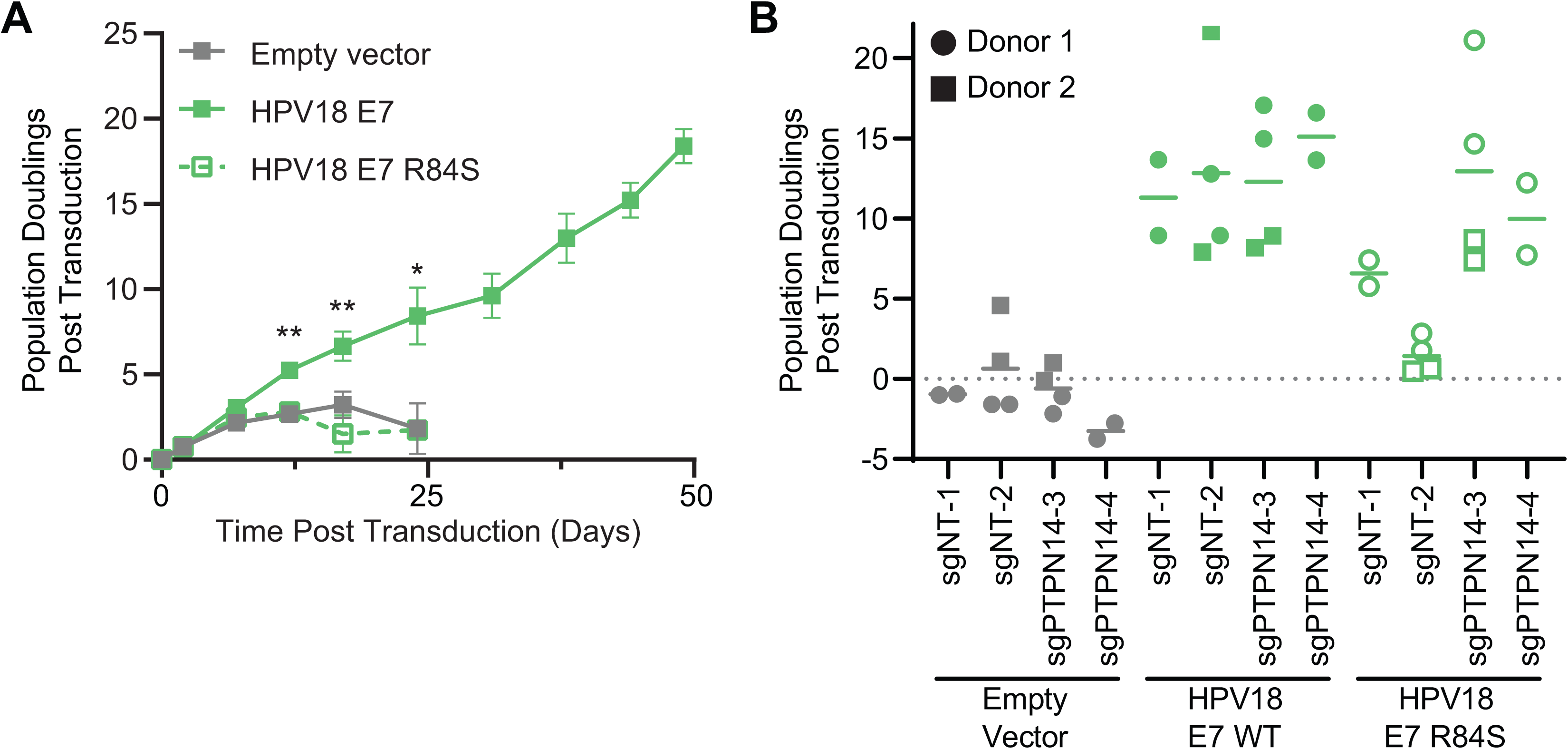
PTPN14 degradation is required for HPV18 E7 to extend the lifespan of primary human keratinocytes. (A) Primary HFK were transduced with retroviruses encoding HPV18 E7 WT, HPV18 E7 R84S, or the empty vector control and passaged for up to 49 days. Graph displays the growth of these cell lines over the course of the experiment. Statistical significance was determined from three independent experiments by repeated measures two-way ANOVA (\**p* < 0.05; \*\**p* < 0.01). (B) Primary HFK from two different donors were transduced with the same retroviruses as *(A)* along with LentiCRISPRv2 lentiviral vectors encoding SpCas9 and nontargeting or PTPN14-directed sgRNAs and cultured for up to 58 days. Graph displays the final population doubling values under each condition along with the mean.

## DISCUSSION

HPV E7 from diverse virus genotypes bind and degrade PTPN14 dependent on the host E3 ubiquitin ligase UBR4 (11, 28, 30, 36). A potential tumor suppressive role of PTPN14 is consistent with its mutation rates in certain cancers (basal cell carcinoma: 23%; gastrointestinal cancers: 5%) (46, 47) and its role as an inhibitor of the HIPPO pathway proto-oncogenes YAP and TAZ (32, 34, 47). The high degree of conservation of E7-PTPN14 binding suggests that PTPN14 binding and/or degradation play an important role in HPV biology. We hypothesize that one consequence of PTPN14 degradation is a contribution to high-risk HPV-mediated carcinogenesis.

High-risk E7 is central to carcinogenesis. In a recent analysis of the HPV16 sequences from >5000 patient samples, E7 was found to be the most highly conserved HPV ORF in high-grade lesions. This was interpreted to reflect strong selective pressure to maintain prototypical E7 sequences during carcinogenic progression. The requirement for UBR4 in PTPN14 degradation connects PTPN14 degradation to HPV-mediated carcinogenesis. Binding to UBR4 was reported by several groups to contribute to the carcinogenic activities of oncogenic papillomavirus E7 (23, 48). We recently characterized the HPV16 E7 E10K variant that cannot bind UBR4 and is unable to degrade PTPN14 (29). We found that HPV16 E7 E10K was impaired in assays of E7 carcinogenic activity. Using HPV16 E7 E10K, we also discovered that HPV16 E7 represses keratinocyte differentiation downstream of PTPN14 degradation.

In the current study we characterized a newly described E7 variant with a mutation in a highly conserved C-terminal arginine residue. A recently published crystal structure of HPV18 E7 and PTPN14 determined that the conserved arginine is a central component of the E7-PTPN14 binding interface (36). We found that mutation of this arginine rendered the E7 proteins from four HPV genotypes across three species unable to degrade PTPN14 (Figure 1). Thus, the PTPN14-binding interface appears to be highly conserved, particularly among high-risk HPV E7 proteins. In contrast, mutation of an acidic amino acid in the E7 N-terminus (the UBR4 binding region) had a somewhat variable effect on PTPN14 levels. The HPV16 E7 E10K mutation almost completely restored PTPN14 protein levels whereas the analogous HPV18 E7 D10K mutation had an intermediate effect. One possibility is that HPV18 E7 D10K retains some capacity to bind to UBR4.

Our prior work showed that the E7/PTPN14/UBR4 protein complex is distinct from the E7/RB1 complex (28) and that the impacts of the two on the keratinocyte transcriptome are genetically separable (29). We confirmed that HPV16 E7 R77S and HPV18 E7 R84S retain the capacity to bind and degrade RB1 (Figures 1 and 3) and promote the expression of E2F target genes (Figures 2 and 6). These data strongly support the model that PTPN14 binding and degradation is a conserved activity of HPV E7 separate from RB1 inactivation. The rare HPV16 E7 R77S and E10K variants were found by Mirabelllo and colleagues in HPV16-positive precancerous lesions (35). We do not know whether genetic polymorphisms or a characteristic of the tumor microenvironment in the patients harboring the variants enabled their progression past the initial infection nor whether the lesions would have progressed to frank carcinoma.

Either PTPN14 degradation by HPV16 E7 or PTPN14 CRIPSR KO represses the expression of differentiation-related genes (29). In this study, we confirmed that blocking PTPN14 binding through mutation of the conserved C-terminal arginine likewise renders high-risk HPV16 E7 and HPV18 E7 unable to repress differentiation (Figures 4 and 5). Furthermore, we confirmed by RNA-seq analysis that repression of differentiation is the primary effect of PTPN14 binding by HPV18 E7 on the transcriptome of unstimulated primary keratinocytes (Figure 6). Other alpha papillomavirus proteins including E6 repress differentiation, although the downstream mechanisms remain to be elucidated (41–43). Mouse PTPN14 is proposed to be a downstream effector of p53 in the “p53-PTPN14-YAP axis” (47). If PTPN14 is also downstream of p53 in human cells, inactivation of PTPN14 by E7 could be more complex in the context of p53 degradation by E6. Nonetheless, we observed substantially higher expression of differentiation marker expression in HFK harboring the HPV18^E7 R84S^ genome than those harboring the HPV18^E7 WT^ genome (Figure 7). This indicates that PTPN14 degradation contributes to repression of differentiation even in the presence of E6. Beta papillomavirus E6 represses differentiation by binding to MAML and directly inhibiting NOTCH signaling (49–52). However, alpha papillomavirus E6 proteins do not bind MAML1 (53, 54). We posit that all papillomaviruses require repression of differentiation gene expression and that alpha HPV depend on PTPN14 degradation by E7 to repress differentiation.

Although HPV E7 repress differentiation-related genes through PTPN14 degradation, our experiments indicate that E7-expressing cells retain the capacity to ultimately differentiate, which is consistent with the differentiation-dependent nature of the HPV life cycle (55–60). Additionally, we observed that PTPN14 binding by HPV18 E7 had a much stronger effect on cell viability after growth in suspension (Figure 5B) than it did on differentiation-related gene expression in pools of differentiated keratinocytes (Figures 4 and 5A). These observations may be related to the stage of the keratinocyte differentiation program at which PTPN14 degradation has the most impact. For example, it is possible that PTPN14 affects cell fate decisions early in differentiation. Our findings are consistent with PTPN14 binding by E7 delaying or impairing the commitment to terminal differentiation—thereby allowing for continued proliferation of some cells—as opposed to completely blocking the differentiation program.

High-risk HPV E7 expression extends the life span of primary keratinocytes (8, 45) and lifespan extension is directly correlated with E7 carcinogenic activity. Using lifespan extension experiments in primary HFK, we found that HPV18 E7 R84S was substantially impaired in its ability to extend the lifespan of primary cells compared to WT HPV18 E7 (Figure 8A). Moreover, CRISPR/Cas9 knockout of PTPN14 in primary HFK rescued the lifespan-extending activity of HPV18 E7 R84S (Figure 8B). HPV E7 can bind to PTPN21 (11, 36), a homolog of PTPN14, but the rescue experiment indicates that PTPN14, not PTPN21, is the primary target involved in lifespan extension. These data and our analysis of survival after growth in suspension (Figure 5B) implicate PTPN14 binding and degradation as a carcinogenic activity of high-risk HPV E7.

Our finding that PTPN14 binding and degradation leads to repression of differentiation-related genes also has implications for its role in carcinogenesis. This activity is significant because it suggests that PTPN14 degradation could contribute to the tendency for HPV-positive oropharyngeal carcinomas to be poorly differentiated (61, 62). It is interesting to consider that targeting PTPN14 binding to promote differentiation could present an opportunity for the treatment of HPV positive cancers. All-trans retinoic acid (ATRA) is a highly successful differentiation therapy used to treat acute promyelocytic leukemia (63). ATRA has shown promise in mouse models of non-melanoma skin cancer and in reducing proliferation of human squamous cell carcinoma cells (64–66).

Here we show that PTPN14 binding and degradation is a highly conserved activity of HPV E7 proteins. By characterizing an HPV E7 variant that cannot bind PTPN14, we have also shown that inactivation of PTPN14 rather than another target of E7 or of UBR4 leads to the repression of differentiation. Furthermore, we found that the PTPN14 degradation-dependent repression of differentiation occurs even when E7 is expressed from the complete HPV18 genome. Finally, we demonstrated that PTPN14 degradation contributes to the carcinogenic properties of HPV18 E7 and confirmed that PTPN14 KO can rescue this activity. Future studies aimed at elucidating the mechanism by which PTPN14 degradation represses differentiation, promotes carcinogenesis, and influences the differentiation-dependent HPV life cycle will further our understanding of this highly conserved HPV E7 activity.

## MATERIALS AND METHODS

### Plasmids and cloning

Mutations in HPV E7 ORFs were introduced by site-directed mutagenesis into pDONR-Kozak-E7 plasmids and recombined into MSCV-P C-FlagHA GAW or MSCV-Neo C-HA GAW as previously described (11). LentiCRISPR v2 (Addgene # 52961) vectors were cloned according to standard protocols using sgRNA sequences as contained in the Broad Institute Brunello library (67). pNeo-loxP-HPV18 was the kind gift of Drs. Thomas Broker and Louise Chow (44) and was used as the template to generate pNeo-loxP-HPV18 E7 R84S by PCR site-directed mutagenesis of the E7-containing portion of the genome, followed by reassembly of this fragment into the pNeo-loxP-HPV18 vector using HiFi DNA Assembly (NEB). Plasmids used in the study are listed in Supplemental Table 1.

### Cells

Primary human foreskin keratinocytes (HFK) from two donors were isolated and provided as de-identified material by the Penn Skin Biology and Diseases Resource-based Center Core B. N/Tert-1 are hTert-immortalized HFK (68). Keratinocytes were cultured as previously described (29). HPV E7 expressing cell populations were established by transduction with MSCV retroviral vectors followed by puromycin selection. PTPN14 knockout or nontargeting control primary HFK were established by transduction with LentiCRISPR v2 vectors followed by puromycin selection. HPV E7 expression in the CRISPR/Cas9-edited cells was achieved by simultaneous transduction of MSCV retroviral vectors and selection with G418 and puromycin. Retroviruses and lentiviruses were generated as previously described (11). To generate HFK containing complete HPV genomes, cells were co-transfected with pNeo-loxP-HPV18 or pNeo-loxP-HPV18 E7 R84S and a Cre expression plasmid using FuGENE HD (Promega) as previously described (44). HFK populations containing the HPV genomes were selected for with G418 for 4 days.

### Western blotting

Western blots were performed using Mini-PROTEAN or Criterion (BioRad) SDS-PAGE gels and transfer to PVDF. After blocking in 5% nonfat dried milk in TBS-T (Tris buffered saline [pH 7.4] with 0.05% Tween-20), membranes were incubated with primary antibodies as follows: RB1 (Calbiochem/EMD), actin (Millipore), PTPN14 (R&D Systems or Cell Signaling Technology), KRT1 (Santa Cruz), KRT10 (Santa Cruz), or involucrin (Santa Cruz)). Membranes were washed in TBS-T and incubated with horseradish peroxidase (HRP)-coupled anti-mouse or anti-rabbit antibodies and detected using chemiluminescent substrate. HA-tagged proteins were detected and visualized using an HA antibody conjugated to HRP (Roche). For anti-HA immunoprecipitations, HA-tagged proteins were immunoprecipitated and processed for Western blot as previously described (28).

### qRT-PCR

Total RNA was isolated from N/Tert or HFK cells using the NucleoSpin RNA extraction kit (Macherey-Nagel). RNA was reverse transcribed using the High Capacity cDNA Reverse Transcription Kit (Applied Biosystems). cDNAs were assayed by qPCR using Fast SYBR Green Master Mix (Applied Biosystems) using a QuantStudio 3 (ThermoFisher). KiCqStart SYBR green primers for qRT-PCR (Sigma-Aldrich) were specific for MCM2, CCNE2, KRT1, KRT10, IVL, MAF, GAPDH, and G6PD. All qRT-PCR data were normalized to GAPDH or to G6PD.

### Keratinocyte Differentiation Assays

N/Tert-E7 or empty vector cells were stimulated to differentiate by growth in high-calcium medium or by culture in suspension. For calcium differentiation, cells were grown to ~30% confluence in 6-well plates using standard growth conditions in Keratinocyte Serum-Free Medium (K-SFM) with growth factors. To induce differentiation, growth factors were withdrawn and cells were refed with Keratinocyte Basal Medium (Lonza) supplemented to achieve 1.5 mM [final] CaCl_2_. Cells were harvested for RNA analysis 72h post-treatment. For culture in suspension, cells were harvested by trypsinization and re-plated in ultra-low attachment plates (Sigma-Aldrich). After 0 or 12h of culture in suspension cells were harvested for RNA analysis or 1000 cells were re-plated in standard 6-well tissue culture plates. Re-plated cells were stained with crystal violet 5 days post re-plating. pNeo-loxP-HPV18 transfected HFK cells were also grown to high confluence and stimulated to differentiate by culturing for 0, 2, or 6 days in growth factor-free Keratinocyte Basal Medium (Lonza) supplemented to achieve 1.5 mM [final] CaCl_2_.

### RNA-seq

Total RNA was isolated from 3 independent isolates of HFK-empty vector control, HFK-18E7, or HFK 18E7 R84S cells using the RNeasy mini kit (Qiagen). PolyA selection, reverse transcription, library construction, sequencing, and initial analysis were performed by Novogene. Differentially expressed genes were selected based on a 1.5-fold change and FDR-adjusted *p* < 0.05 cutoff and were analyzed for enriched biological processes (BP) using the GO enrichment analysis tool of the PANTHER classification system (69). All GO terms in enrichment analyses are displayed in rank order by FDR-adjusted *p*-value. RNA-seq data have been deposited in NCBI GEO with accession number GSE150201.

### Keratinocyte Lifespan Extension

To assess the ability of HPV18 E7 variants to extend keratinocyte lifespan, primary HFK were transduced with MSCV-E7 retroviruses and selected with puromycin. Cells were selected in puromycin and passaged for approximately 50 days. Population doublings were calculated based upon the number of cells collected and re-plated at each passage. Each experiment consisted of three independent replicates conducted simultaneously.

### Data Availability

RNA-seq data from HFK-empty, HFK-18E7, and HFK-18E7 R84S are freely accessible in NCBI GEO with accession number GSE150201.

## Acknowledgements

We thank the members of our laboratories for helpful discussions. This work was supported by American Cancer Society grant 131661-RSG-18-048-01-MPC and NIH R01 AI148431 to EAW and NIH R01 CA066980 to KM. JH was supported by NIH T32 AI007324. The Penn Skin Biology and Diseases Resource-based Center was supported by NIH P30 AR068589.

## REFERENCES

1. de Martel C, Plummer M, Vignat J, Franceschi S. 2017. Worldwide burden of cancer attributable to HPV by site, country and HPV type. Int J Cancer 141:664–670.

2. Graham S, Graham, V. S. 2017. Keratinocyte Differentiation-Dependent Human Papillomavirus Gene Regulation. Viruses 9:245.

3. Van Doorslaer K, Tan Q, Xirasagar S, Bandaru S, Gopalan V, Mohamoud Y, Huyen Y, McBride AA. 2013. The Papillomavirus Episteme: A central resource for papillomavirus sequence data and analysis. Nucleic Acids Res 41:D571–D578.

4. Münger K, Phelps WC, Bubb V, Howley PM, Schlegel R. 1989. The E6 and E7 genes of the human papillomavirus type 16 together are necessary and sufficient for transformation of primary human keratinocytes. J Virol 63:4417–4421.

5. Bedell MA, Jones KH, Grossman SR, Laimins LA. 1989. Identification of human papillomavirus type 18 transforming genes in immortalized and primary cells. J Virol 63:1247–1255.

6. Francis DA, Schmid SI, Howley PM. 2000. Repression of the Integrated Papillomavirus E6/E7 Promoter Is Required for Growth Suppression of Cervical Cancer Cells. J Virol 74:2679–2686.

7. Goodwin EC, DiMaio D. 2000. Repression of human papillomavirus oncogenes in HeLa cervical carcinoma cells causes the orderly reactivation of dormant tumor suppressor pathways. Proc Natl Acad Sci U S A 97:12513–12518.

8. Hawley-Nelson P, Vousden KH, Hubbert NL, Lowy DR, Schiller JT. 1989. HPV16 E6 and E7 proteins cooperate to immortalize human foreskin keratinocytes. EMBO J 8:3905–3910.

9. Dyson N, Guida P, Münger K, Harlow E. 1992. Homologous sequences in adenovirus E1A and human papillomavirus E7 proteins mediate interaction with the same set of cellular proteins. J Virol 66:6893–902.

10. Münger K, Werness BA, Dyson N, Phelps WC, Harlow E, Howley PM. 1989. Complex formation of human papillomavirus E7 proteins with the retinoblastoma tumor suppressor gene product. EMBO J 8:4099–4105.

11. White EA, Sowa ME, Tan MJA, Jeudy S, Hayes SD, Santha S, Münger K, Harper JW, Howley PM. 2012. Systematic identification of interactions between host cell proteins and E7 oncoproteins from diverse human papillomaviruses. Proc Natl Acad Sci U S A 109:E260–7.

12. Huh K, Zhou X, Hayakawa H, Cho J-Y, Libermann TA, Jin J, Harper JW, Munger K. 2007. Human papillomavirus type 16 E7 oncoprotein associates with the cullin 2 ubiquitin ligase complex, which contributes to degradation of the retinoblastoma tumor suppressor. J Virol 81:9737–47.

13. Scheffner M, Werness BA, Huibregtse JM, Levine AJ, Howley PM. 1990. The E6 oncoprotein encoded by human papillomavirus types 16 and 18 promotes the degradation of p53. Cell 63:1129–1136.

14. Werness BA, Levine AJ, Howley PM. 1990. Association of human papillomavirus types 16 and 18 E6 proteins with p53. Science 248:76–79.

15. Sharma S, Munger K. 2020. KDM6A mediated expression of the long noncoding RNA DINO causes TP53 tumor suppressor stabilization in Human Papillomavirus type 16 E7 expressing cells. J Virol 02178–19.

16. Seavey SE, Holubar M, Saucedo LJ, Perry ME. 1999. The E7 Oncoprotein of Human Papillomavirus Type 16 Stabilizes p53 through a Mechanism Independent of p19ARF. J Virol 73:7590–7598.

17. Ciccolini F, Di Pasquale G, Carlotti F, Crawford L, Tommasino M. 1994. Functional studies of E7 proteins from different HPV types. Oncogene 9:2633–8.

18. Ibaraki T, Satake M, Kurai N, Ichijo M, Ito Y. 1993. Transacting activities of the E7 genes of several types of human papillomavirus. Virus Genes 7:187–196.

19. Balsitis S, Dick F, Lee D, Farrell L, Hyde RK, Griep AE, Dyson N, Lambert PF. 2005. Examination of the pRb-Dependent and pRb-Independent Functions of E7 In Vivo. J Virol 79:11392–11402.

20. Jewers RJ, Hildebrandt P, Ludlow JW, Kell B, McCance DJ. 1992. Regions of human papillomavirus type 16 E7 oncoprotein required for immortalization of human keratinocytes. J Virol 66:1329–35.

21. White EA, Kramer RE, Hwang JH, Pores Fernando AT, Naetar N, Hahn WC, Roberts TM, Schaffhausen BS, Livingston DM, Howley PM. 2015. Papillomavirus E7 Oncoproteins Share Functions with Polyomavirus Small T Antigens. J Virol 89:2857–2865.

22. Banks L, Edmonds C, Vousden KH. 1990. Ability of the HPV16 E7 protein to bind RB and induce DNA synthesis is not sufficient for efficient transforming activity in NIH3T3 cells. Oncogene 5:1383–9.

23. Huh KW, DeMasi J, Ogawa H, Nakatani Y, Howley PM, Münger K. 2005. Association of the human papillomavirus type 16 E7 oncoprotein with the 600-kDa retinoblastoma protein-associated factor, p600. Proc Natl Acad Sci U S A 102:11492–11497.

24. Strati K, Lambert PF. 2007. Role of Rb-dependent and Rb-independent functions of papillomavirus E7 oncogene in head and neck cancer. Cancer Res 67:11585–11593.

25. Balsitis S, Dick F, Dyson N, Lambert PF. 2006. Critical Roles for Non-pRb Targets of Human Papillomavirus Type 16 E7 in Cervical Carcinogenesis. Cancer Res 66:9393–400.

26. Phelps WC, Münger K, Yee CL, Barnes JA, Howley PM. 1992. Structure-function analysis of the human papillomavirus type 16 E7 oncoprotein. J Virol 66:2418–2427.

27. Helt A-MA-M, Galloway DA. 2002. Destabilization of the Retinoblastoma Tumor Suppressor by Human Papillomavirus Type 16 E7 Is Not Sufficient To Overcome Cell Cycle Arrest in Human Keratinocytes. J Virol 75:6737–6747.

28. White EA, Münger K, Howley PM. 2016. High-Risk Human Papillomavirus E7 Proteins Target PTP N14 for Degradation. MBio 7:e01530–16.

29. Hatterschide J, Bohidar AE, Grace M, Nulton TJ, Kim HW, Windle B, Morgan IM, Munger K, White EA. 2019. PTPN14 degradation by high-risk human papillomavirus E7 limits keratinocyte differentiation and contributes to HPV-mediated oncogenesis. Proc Natl Acad Sci U S A 116:7033–7042.

30. Szalmás A, Tomaić V, Basukala O, Massimi P, Mittal S, Kónya J, Banks L. 2017. The PTPN14 Tumor Suppressor Is a Degradation Target of Human Papillomavirus E7. J Virol 91:e00057–17.

31. Wilson KE, Li Y-W, Yang N, Shen H, Orillion AR, Zhang J. 2014. PTPN14 forms a complex with Kibra and LATS1 proteins and negatively regulates the YAP oncogenic function. J Biol Chem 289:23693–700.

32. Knight JF, Sung VYC, Kuzmin E, Couzens AL, de Verteuil DA, Ratcliffe CDH, Coelho PP, Johnson RM, Samavarchi-Tehrani P, Gruosso T, Smith HW, Lee W, Saleh SM, Zuo D, Zhao H, Guiot MC, Davis RR, Gregg JP, Moraes C, Gingras AC, Park M. 2018. KIBRA (WWC1) Is a Metastasis Suppressor Gene Affected by Chromosome 5q Loss in Triple-Negative Breast Cancer. Cell Rep 22:3191–3205.

33. Lin Z, Yang Z, Xie R, Ji Z, Guan K, Zhang M. 2019. Decoding WW domain tandem-mediated target recognitions in tissue growth and cell polarity. Elife 8:e49439.

34. Wang W, Huang J, Wang X, Yuan J, Li X, Feng L, Park J Il, Chen J. 2012. PTPN14 is required for the density-dependent control of YAP1. Genes Dev 26:1959–1971.

35. Mirabello L, Yeager M, Yu K, Clifford GM, Xiao Y, Zhu B, Cullen M, Boland JF, Wentzensen N, Nelson CW, Raine-Bennett T, Chen Z, Bass S, Song L, Yang Q, Steinberg M, Burdett L, Dean M, Roberson D, Mitchell J, Lorey T, Franceschi S, Castle PE, Walker J, Zuna R, Kreimer AR, Beachler DC, Hildesheim A, Gonzalez P, Porras C, Burk RD, Schiffman M. 2017. HPV16 E7 Genetic Conservation Is Critical to Carcinogenesis. Cell 170:1164–1174.e6.

36. Yun H-Y, Kim MW, Lee HS, Kim W, Shin JH, Kim H, Shin H-C, Park H, Oh B-H, Kim WK, Bae K-H, Lee SC, Lee E-W, Ku B, Kim SJ. 2019. Structural basis for recognition of the tumor suppressor protein PTPN14 by the oncoprotein E7 of human papillomavirus. PLOS Biol 17:e3000367.

37. Adams JC, Watt FM. 1989. Fibronectin inhibits the terminal differentiation of human keratinocytes. Nature 340:307–309.

38. Banno T, Blumenberg M. 2014. Keratinocyte detachment-differentiation connection revisited, or Anoikis-Pityriasi Nexus Redux. PLoS One 9:1–12.

39. Green H. 1977. Terminal differentiation of cultured human epidermal cells. Cell 11:405–416.

40. Lopez-Pajares V, Qu K, Zhang J, Webster DE, Barajas BC, Siprashvili Z, Zarnegar BJ, Boxer LD, Rios EJ, Tao S, Kretz M, Khavari PA. 2015. A LncRNA-MAF:MAFB transcription factor network regulates epidermal differentiation. Dev Cell 32:693–706.

41. Duffy CL, Phillips SL, Klingelhutz AJ. 2003. Microarray analysis identifies differentiation-associated genes regulated by human papillomavirus type 16 E6. Virology 314:196–205.

42. Zehbe I, Richard C, DeCarlo CA, Shai A, Lambert PF, Lichtig H, Tommasino M, Sherman L. 2008. Human papillomavirus 16 E6 variants differ in their dysregulation of human keratinocyte differentiation and apoptosis. Virology 383:69–77.

43. White EA. 2019. Manipulation of Epithelial Differentiation by HPV Oncoproteins. Viruses 1–25.

44. Wang H-K, Duffy AA, Broker TR, Chow LT. 2009. Robust production and passaging of infectious HPV in squamous epithelium of primary human keratinocytes. Genes Dev 23:181–94.

45. Halbert CL, Demers GW, Galloway DA. 1991. The E7 gene of human papillomavirus type 16 is sufficient for immortalization of human epithelial cells. J Virol 65:473–8.

46. Bonilla X, Parmentier L, King B, Bezrukov F, Kaya G, Zoete V, Seplyarskiy VB, Sharpe HJ, McKee T, Letourneau A, Ribaux PG, Popadin K, Basset-Seguin N, Chaabene R Ben, Santoni FA, Andrianova MA, Guipponi M, Garieri M, Verdan C, Grosdemange K, Sumara O, Eilers M, Aifantis I, Michielin O, de Sauvage FJ, Antonarakis SE, Nikolaev SI. 2016. Genomic analysis identifies new drivers and progression pathways in skin basal cell carcinoma. Nat Genet 48:398–406.

47. Mello SS, Valente LJ, Raj N, Seoane JA, Flowers BM, McClendon J, Bieging-Rolett KT, Lee J, Ivanochko D, Kozak MM, Chang DT, Longacre TA, Koong AC, Arrowsmith CH, Kim SK, Vogel H, Wood LD, Hruban RH, Curtis C, Attardi LD. 2017. A p53 Super-tumor Suppressor Reveals a Tumor Suppressive p53-Ptpn14-Yap Axis in Pancreatic Cancer. Cancer Cell 32:460–473.e6.

48. DeMasi J, Huh KW, Nakatani Y, Münger K, Howley PM. 2005. Bovine papillomavirus E7 transformation function correlates with cellular p600 protein binding. Proc Natl Acad Sci U S A 102:11486–11491.

49. Meyers JM, Uberoi A, Grace M, Lambert PF, Munger K. 2017. Cutaneous HPV8 and MmuPV1 E6 Proteins Target the NOTCH and TGF-β Tumor Suppressors to Inhibit Differentiation and Sustain Keratinocyte Proliferation. PLOS Pathog 13:e1006171.

50. Meyers JM, Spangle JM, Munger K. 2013. The Human Papillomavirus Type 8 E6 Protein Interferes with NOTCH Activation during Keratinocyte Differentiation. J Virol 87:4762–4767.

51. Brimer N, Lyons C, Wallberg AE, Vande Pol SB. 2012. Cutaneous papillomavirus E6 oncoproteins associate with MAML1 to repress transactivation and NOTCH signaling. Oncogene 31:4639–4646.

52. Tan MJA, White EA, Sowa ME, Harper JW, Aster JC, Howley PM. 2012. Cutaneous β-human papillomavirus E6 proteins bind Mastermind-like coactivators and repress Notch signaling. Proc Natl Acad Sci U S A 109:E1473–E1480.

53. White EA, Kramer RE, Tan MJA, Hayes SD, Harper JW, Howley PM. 2012. Comprehensive Analysis of Host Cellular Interactions with Human Papillomavirus E6 Proteins Identifies New E6 Binding Partners and Reflects Viral Diversity. J Virol 86:13174–13186.

54. Rozenblatt-Rosen O, Deo RC, Padi M, Adelmant G, Calderwood MA, Rolland T, Grace M, Dricot A, Askenazi M, Tavares M, Pevzner SJ, Abderazzaq F, Byrdsong D, Carvunis AR, Chen AA, Cheng J, Correll M, Duarte M, Fan C, Feltkamp MC, Ficarro SB, Franchi R, Garg BK, Gulbahce N, Hao T, Holthaus AM, James R, Korkhin A, Litovchick L, Mar JC, Pak TR, Rabello S, Rubio R, Shen Y, Singh S, Spangle JM, Tasan M, Wanamaker S, Webber JT, Roecklein-Canfield J, Johannsen E, Barabási AL, Beroukhim R, Kieff E, Cusick ME, Hill DE, Münger K, Marto JA, Quackenbush J, Roth FP, Decaprio JA, Vidal M. 2012. Interpreting cancer genomes using systematic host network perturbations by tumour virus proteins. Nature 487:491–495.

55. Graham S V. 2017. The human papillomavirus replication cycle, and its links to cancer progression: a comprehensive review. Clin Sci 131:2201–2221.

56. Doorbar J, Egawa N, Griffin H, Kranjec C, Murakami I. 2015. Human papillomavirus molecular biology and disease association. Rev Med Virol 25:2–23.

57. McBride AA. 2017. Mechanisms and strategies of papillomavirus replication. Biol Chem 398:919–927.

58. Peh WL, Middleton K, Christensen N, Nicholls P, Egawa K, Sotlar K, Brandsma J, Percival A, Lewis J, Liu WJ, Doorbar J. 2002. Life cycle heterogeneity in animal models of human papillomavirus-associated disease. J Virol 76:10401–16.

59. Moody CA, Fradet-Turcotte A, Archambault J, Laimins LA. 2007. Human papillomaviruses activate caspases upon epithelial differentiation to induce viral genome amplification. Proc Natl Acad Sci 104:19541–19546.

60. Pentland I, Campos-León K, Cotic M, Davies K-J, Wood CD, Groves IJ, Burley M, Coleman N, Stockton JD, Noyvert B, Beggs AD, West MJ, Roberts S, Parish JL. 2018. Disruption of CTCF-YY1–dependent looping of the human papillomavirus genome activates differentiation-induced viral oncogene transcription. PLOS Biol 16:e2005752.

61. Pai SI, Westra WH. 2009. Molecular Pathology of Head and Neck Cancer: Implications for Diagnosis, Prognosis, and Treatment. Annu Rev Pathol Mech Dis 4:49–70.

62. Mendelsohn AH, Lai CK, Shintaku IP, Elashoff DA, Dubinett SM, Abemayor E, St. John MA. 2010. Histopathologic findings of HPV and p16 positive HNSCC. Laryngoscope 120:1788–1794.

63. Ablain J, De Thé H. 2014. Retinoic acid signaling in cancer: The parable of acute promyelocytic leukemia. Int J Cancer 135:2262–2272.

64. Zhang ML, Tao Y, Zhou WQ, Ma PC, Cao YP, He CD, Wei J, Li LJ. 2014. All-trans retinoic acid induces cell-cycle arrest in human cutaneous squamous carcinoma cells by inhibiting the mitogen-activated protein kinase-activated protein 1 pathway. Clin Exp Dermatol 39:354–360.

65. Verma AK. 1987. Inhibition of Both Stage I and Stage II Mouse Skin Tumor Promotion by Retinoic Acid and the Dependence of Inhibition of Tumor Promotion on the Duration of Retinoic Acid Treatment. Cancer Res 47:5092–5140.

66. Cheepala SB, Yin W, Syed Z, Gill JN, McMillian A, Kleiner HE, Lynch M, Loganantharaj R, Trutschl M, Cvek U, Clifford JL. 2009. Identification of the B-Raf/Mek/Erk MAP kinase pathway as a target for all-trans retinoic acid during skin cancer promotion. Mol Cancer 8:27.

67. Doench JG, Fusi N, Sullender M, Hegde M, Vaimberg EW, Donovan KF, Smith I, Tothova Z, Wilen C, Orchard R, Virgin HW, Listgarten J, Root DE. 2016. Optimized sgRNA design to maximize activity and minimize off-target effects of CRISPR-Cas9. Nat Biotechnol 34:184–191.

68. Dickson MA, Hahn WC, Ino Y, Ronfard V, Wu JY, Weinberg RA, Louis DN, Li FP, Rheinwald JG. 2000. Human Keratinocytes That Express hTERT and Also Bypass a p16INK4a-Enforced Mechanism That Limits Life Span Become Immortal yet Retain Normal Growth and Differentiation Characteristics. Mol Cell Biol 20:1436–1447.

69. Mi H, Huang X, Muruganujan A, Tang H, Mills C, Kang D, Thomas PD. 2017. PANTHER version 11: Expanded annotation data from Gene Ontology and Reactome pathways, and data analysis tool enhancements. Nucleic Acids Res 45:D183–D189.

